# Human life expectancies are still rising

**DOI:** 10.1101/2025.05.01.651310

**Authors:** Lucio Vinicius, Andrea Migliano

## Abstract

Olshansky and collaborators have recently proposed that the era of continuous extension of human lifespans has finally come to an end^1^. By analysing thirty years of recent demographic data (1990-2020) from ten populations (the eight longest-lived nations, plus Hong Kong and the USA), they rejected the claims made by Oeppen and Vaupel at the start of the century^2^ (and more recently by Vaupel and collaborators^3^) that human longevity was still far from approaching an upper limit. However, on closer examination the results by Olshansky and colleagues seem to complement rather than directly challenge the radical life extension hypothesis. The reason is that the latter was based not on country-level demographic patterns but instead on a best-practice life expectancy trend resulting from the succession of annual world-leading populations. Here we present an update based on data from the last two decades that confirms Oeppen and Vaupel’s original insights and demonstrates that both female and male lifespans continue to linearly increase at a global scale. This remarkably long trend observed since 1840 remains at odds with our expectation that human lifespans must at some point hit a biologically imposed ceiling.

## MAIN TEXT

Olshansky et al. detected recent changes in key biodemographic parameters in each of their sampled populations, such as overall reductions in annual rates of lifespan extension and decreases in lifespan inequality, pointing to a compression in age at death distributions and vanishing potential for further longevity increases. However, these and other country-level analyses do not capture the fact that the world-leading population in female longevity changed over 40 times between 1840 and 2001 in Oeppen’s and Vaupel’s original series. Figure 1 extends their original best-practice regression line by incorporating new data from 2001 to 2020 from the Human Mortality Database^4^, although some data are available until 2023 (to avoid the effects of the COVID-19 pandemics), as well as data for men. The updated series shows that in the first two decades of this century female maximum lifespans have again increased from 84.85 to 87.75 years, a rate of 1.45 years per decade, lower than the 2.49 years in the previous two decades, and 2.31 years per decade since 1840. In men, the rate of increase was 1.96 years per decade in this century, very similar to the values of 2.02 in the previous two decades and 2.03 per decade since 1840. Despite the slower improvement in women, data from this century sit comfortably within prediction bands of the general regression, with the recent rates per year or decade not being the lowest since 1840 or (1940) either for women or men (Figures S2-S5). Since all previous claims (beginning in the 1940s) that humans were approaching a longevity ceiling have been shown to be premature, it seems that data from the last two decades do not yet provide conclusive evidence that the long-term global trend has come to a halt.

**Figure 1.**
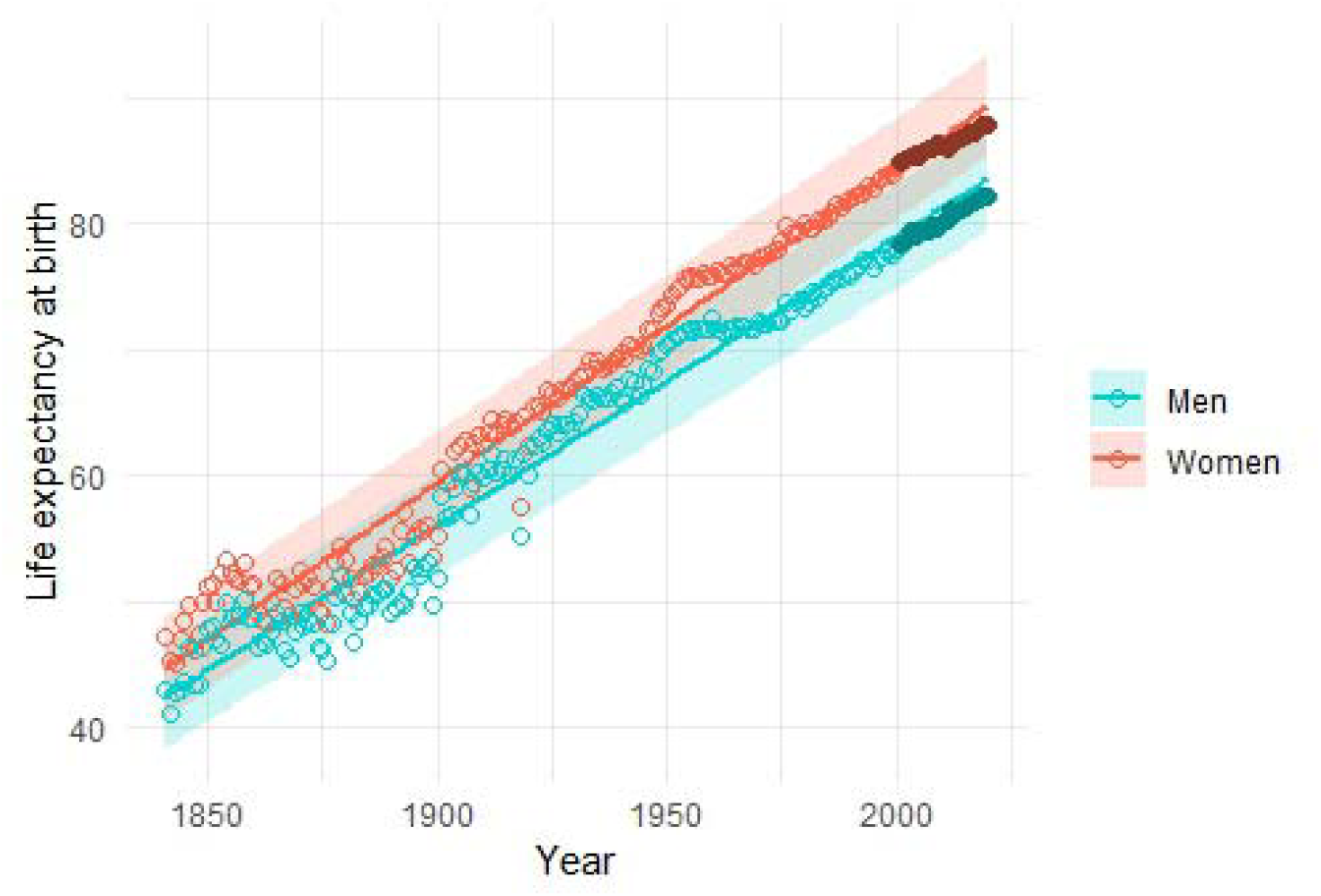
Life expectancy in the world-leading population, 1840-2020. Solid circles are data from 2001-2020 (see refs 1 and 2 for list of nations). Regression slopes are b = 0.249 (R^2^ = 0.977) for women, and b = 0.229 (R^2^ = 0.971). See Figs. S6 and S7 for versions of this figure including the names of the world-leading populations.

Although the country-level findings by Olshansky and colleagues are solid and of utmost importance for policy making, they do not directly address the best-practice trend described above. Their attempt to draw general conclusions about the limits to human longevity from ‘the fundamental relation between life expectancy and the demographic metrics of life table entropy and lifespan inequality’ is therefore questionable, since such relation was also derived at country level. They also revealed that ‘the most recent decade of change in life expectancy is slower than it was in the last decade of the twentieth century’ in the selected populations. However, such declines at country level are in fact not limited to the most recent decade, and had been identified by Oeppen and Vaupel as occurring in many long-lived populations since at least the 1940s (in parallel to global-level increases in life expectancies). Perhaps for this reason their predicted maximum human lifespans derived from country-level metrics also do not seem to be confirmed by available data. In a previous study^5^, Olshansky and colleagues predicted that in this century ‘*e*_(0)_ for national populations would not likely exceed approximately 85□years (88 for females and 82 for males) unless an intervention in biological aging was discovered’^1^. However, in 2020 male life expectancy in Hong Kong was already 82.17 years (and despite the effects of COVID-2019, in 2023 overall life expectancy was 85.25 years, female life expectancy 87.98 years, and male life expectancy 82.48 years in Hong Kong; in 2023 male life expectancy was also 82.21 years in Switzerland)^4^.

Given the above, we are left with the task of reconciling the signs of a continuing positive trend at global level with the solid evidence for decreasing rates of improvement at country level presented by Olshansky and collaborators. First, the radical lifespan extension hypothesis acknowledged that human lifespans were limited, since ‘modest annual increments in life expectancy will never lead to immortality’^2^. Oeppen and Vaupel’s main question was whether biodemographic evidence was already suggesting convergence towards a longevity ceiling, beyond which improvements in medical care and overall living conditions would no longer have an effect on life expectancies. Over twenty years later, the updated best-practice life expectancy seems to indicate that women and men are not approaching such a limit yet. This is compatible with decreasing rates at country level, since the best-practice trend is the outcome of ‘former laggards catching up with and successively overtaking leading nations’^2^. The underlying logic is straightforward and applies to countless contexts. The energy efficiency of combustion engines, processing speed of microchips, or even the standard of professional tennis underwent undeniable progress over the last decades; nonetheless, the world’s best car engine manufacturer, microchip producer, or professional tennis player have successively changed. By the same token, successive advances in medicine and social welfare may not occur in the same country, with Olshansky and colleagues themselves attributing the recent rise of Hong Kong in the longevity rankings to their recent economic prosperity and tight tobacco control. It remains to be seen whether the link between the best-practice trend and the succession of world leaders may partially explain why men (with six changes and three leading nations; Iceland, Switzerland and Hong Kong) have made faster progress than women (four changes between two populations; Japan and Hong Kong) during the last two decades.

In summary, Olshansky and colleagues have presented undeniable evidence that life expectancies are very unlikely to continue to increase by 2-3 years per decade in each of the longest-lived populations. However, Oeppens and Vaupels’s original insights based on the best-practice trend have not yet been definitely contradicted by biodemographic evidence and remain intriguing. Their original study predicted that female ‘record life expectancy will reach 100 in six decades’ in the 2060s. Over twenty years later, the updated regression above predicts that this will occur in 2063. This remains a credible forecast until there is clearer evidence that the global best-practice life expectancy line is definitely levelling off.

## METHODS

### Data

All raw data used on the text are available on the public Human Mortality Database (https://www.mortality.org/) with longevity data from 41 countries. From the original data, a best-practice dataset was created, consisting of the population with highest female and the highest male longevity for each year between 1841-2023, subsequently filtered to 1841-2020 to avoid the effects of the COVID-19 pandemic.

### Analysis

The extended best-practice curve was then created using *R* and *ggplot2*. Rates of change in longevity per year for women and men were estimated by subtracting consecutive annual longevity values (between 2020 and 1841) from the best-practice dataset. Rates per decade were estimated by subtracting the values at the end of each decade (from 1940 until 2020). All data and code required to replicate the results presented in the text are freely available on https://github.com/LucioUZH/HumanLongevity.

## REFERENCES

1. Olshansky, S.J., Willcox, B.J., Demetrius, L. et al. Implausibility of radical life extension in humans in the twenty-first century. Nat Aging 4, 1635–1642 (2024). 10.1038/s43587-024-00702-3

2. Oeppen, J. & Vaupel, J. W. Broken limits to life expectancy. Science 296, 1029–1031 (2002). 10.1126/science.1069675

3. Vaupel, J. W., Villavicencio, F. & Bergeron□Boucher, M. Demographic perspectives on the rise of longevity. Proc. Natl. Acad. Sci. USA 118, e2019536118 (2021).

4. Human Mortality Database (MPIDR, accessed 21 April 2025); https://www.mortality.org/

5. Olshansky, S. J., Carnes, B. A. & Cassel, C. In search of Methuselah: estimating the upper limits to human longevity. Science 250, 634–640 (1990). 10.1126/science.2237414

