## Supplementary Figures for "Human life expectancies are still rising"

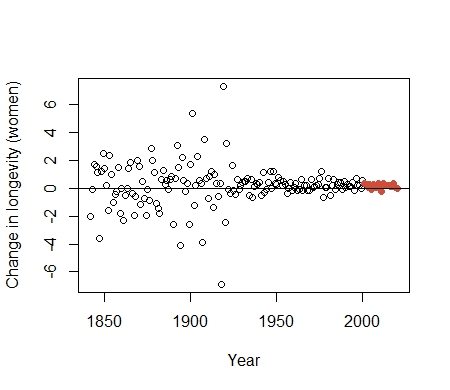

**Figure S1**. Annual change in life expectancy in women since 1840. Red points: data from 2001-2020.

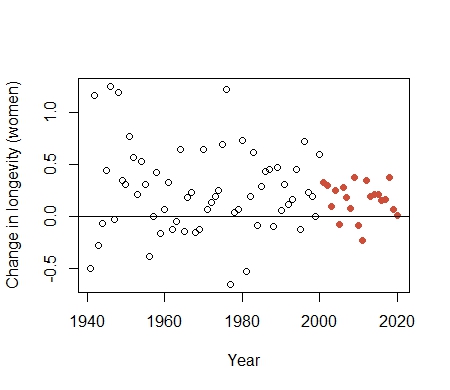

**Figure S2**. Annual change in life expectancy in women since 1940. Red points: data from 2001-2020.

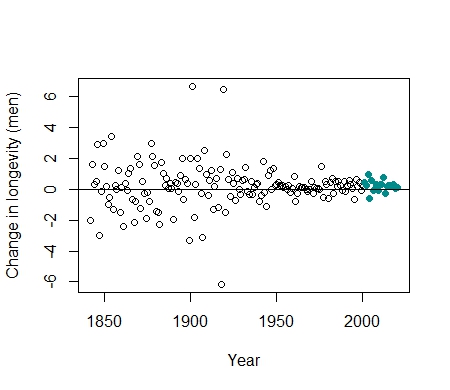

**Figure S3**. Annual change in life expectancy in men since 1840. Green points: data from 2001-2020.

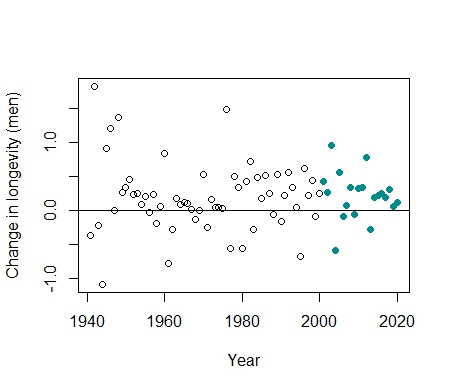

**Figure S4**. Annual change in life expectancy in men since 1940. Green points: data from 2001-2020.

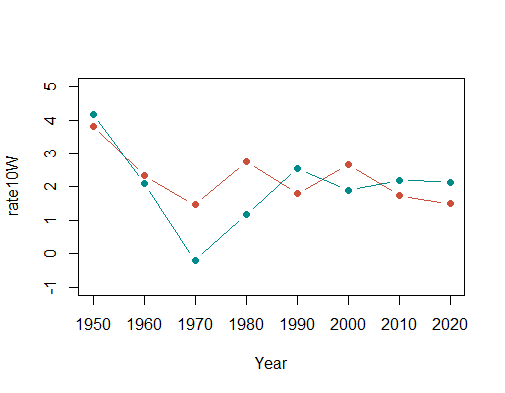

**Figure S5**. Change in life expectancy per decade in women (red) and men (green) since 1950.

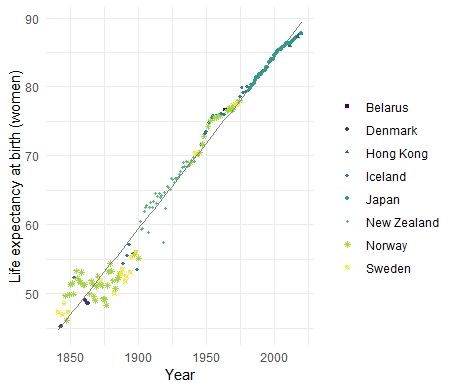

**Figure S6**. Female life expectancy 1840-2020, including key to the world-leading population on each year. Same data points and regression line as in Figure 1.

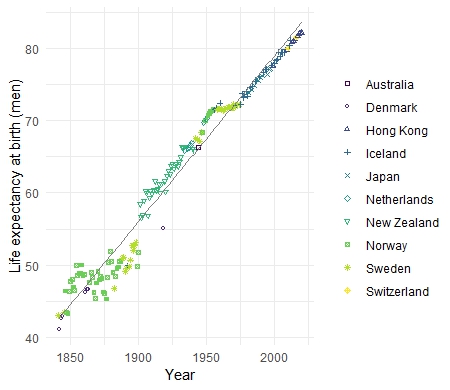

**Figure S7**. Male life expectancy 1840-2020, including key to the world-leading population on each year. Same data points and regression line as in Figure 1.
